# *In silico* and *In vitro* evaluation of the anti-inflammatory and antioxidant potential of *Cymbopogon citratus* from North-western Himalayas

**DOI:** 10.1101/2020.05.31.124982

**Authors:** Deeksha Salaria, Rajan Rolta, Nitin Sharma, Kamal Dev, Anuradha Sourirajan, Vikas Kumar

## Abstract

*Cymbopogon citratus* which is an aromatic perennial herb belonging to family Gramineae is known for its application in food and healthcare industry. The present study attempts to evaluate the potential of essential oil from *Cymbopogon citratus* (CEO) as an anti-inflammatory and antioxidant agent. CEO showed significant DPPH radical scavenging activity (IC_50_ - 91.0 ± 9.25 µg/ml), as compared to Ascorbic acid (IC_50_-33.38 ± 2.29 µg/ml). CEO also exhibited significant *in-vitro* anti-inflammatory activity with IC_50_ - 397.11± 1.45µg/ml) as compared to diclofenac sodium (IC_50_ - 682.98 ± 7.47 µg/ml). Chemical constituents of the oil was determined using Gas Chromatography/Mass Spectroscopy, showed that 8-methyl-3,7-Nonadien-2-one (E), α-Pinene, limonene, citral, limonene oxide and Epoxy-α-terpenyl acetate were the major constituents. The *in silico* molecular docking study showed phytocompounds of CEO (Caryophyllene oxide and β-caryophyllene) have considerable binding potential with 1HD2 and 5IKQ receptors. PASS prediction of these phytocompounds also confirmed strong anti-inflammatory activity of *C. citratus*. The ADMET analysis also showed that these phytocompounds are safer to replace the synthetic drugs with side effects. This work establishes the anti inflammatory potential of CEO as an alternative to existing therapeutic approach to treatment of inflammation and also natural source of antioxidant compounds.

## Introduction

Aromatic and medicinal plants form the backbone of healthcare system for curing various ailments in developing countries including India. Essential oils are regarded as volatile plant components responsible for its aromatic nature [1]. The composition of essential oil varies from plant to plant; but flowers and aerial parts showed comparatively higher amount due to large number of oil producing glands. Because of its strong and pungent aroma, these essential oils primarily serve as insect repellent, thereby protecting them for insects. Other applications of essential oil are as flavoring agent in food and drug industry, starting material for the synthesis of complex medicinal compounds, therapeutic agent for skin and upper respiratory diseases, lipophilic solubility enhancer, carrier of drugs, cosmetic and in fragrance industries, etc [2].

*Cymbopogon citratus* (Gramineae), popularly known as citronella grass or lemongrass is perennial aromatic herb. The pharmacological activities of *C. citratus* have outstanding record in the folk and Ayurvedic medicine [3], [4]. The leaves of lemongrass can be used in both health and food field, as it contains phenol compounds which acts as antioxidant. *C. citratus* has been used for medical purposes to treat pathogenesis neurological disorders. C. citratus is considered as an effective agent in the prevention of various neurological diseases associated with oxidative stress [5]. It is reported to possess antibacterial [6], antifungal [7], antiprotozoal, anti-carcinogenic, anti-inflammatory [8], antioxidant [9], cardioprotective [10], anti-tussive, antiseptic, and anti-rheumatic activities [11]. It has also been used to inhibit platelet aggregation [12], treat diabetes [13], dyslipidemia, gastrointestinal disturbances [7], anxiety, malaria [14], flu, fever, and pneumonia [15], as well as in aromatherapy. In addition to its therapeutic uses, *C. citratus* is also consumed as a tea, added to non-alcoholic beverages, preservative in beverages, baked foods and cuisines [16]. In cosmetics, essential oil of *C. citratus* is used as fragrance in the manufacture of perfumes, soaps, detergents, and creams.

The present work attempts to identify major bioactive molecules present in the essential oil of *Cymbopogon citratus* leaves (CEO) through GC-MS technique. *In vitro* experiments were performed to evaluate antioxidant and anti-inflammatory potential of CEO leaves; Compounds identified through GC-MS were subjected to *in silico* studies to understand the mechanism of antioxidant and anti-inflammatory action. The docking studies predicted that the constituent molecules of *C. citratus* possess more capability as inhibitors as compared to established drugs in the pharmaceutical industry.

## MATERIALS AND METHOD

### Chemicals

The chemicals such as 2,2-diphenyl-1-picrylhydrazyl (DPPH), 2,4,6-Tri(2-pyridyl)-s-triazine (TPTZ), 2,2′-Azino-bis (3-ethylbenzothiazoline-6-sulfonic acid) diammonium salt (ABTS), were obtained from Sigma-Aldrich Co. LLC, Mumbai. Methanol was procured from Loba Chemie Pvt. Ltd., Mumbai. Sodium Diclofenac (100 mg, Ranbaxy Laboratories Mohali, India) and fresh egg albumin were acquired from local market of Solan.

### Collection of *C. citratus* and Extraction of *C. citratus* essential oil (CEO)

*C. citratus* leaves were collected from Palampur, Himachal Pradesh, India (32.1109° N, 76.5363° E) in the month of October 2019. The collected leaves of *C. citratus* were thoroughly washed with distilled water to remove the dust particles. Extraction of CEO was carried out by hydro-distillation method using clevenger assembly [17]. Extraction yield of CEO was determined based on the weight of leaves and oil obtained. The collected CEO was stored at 4 °C for further analysis.

### Gas Chromatography-Mass Spectrometry analysis of CEO

The analysis of essential oil was performed using GC/MS instrument (Thermo Trace 1300 GC coupled with Thermo TSQ 800 Triple Quadrupole MS) fitted with a TG 5MS capillary column (30 m × 0.25 mm, 0.25 µm film thickness). Injector temperature was 280 and 250°C, respectively. The column temperature was held at 45 °C for 8 min and then increased to 250 °C at a rate of 28 °C /min and held at 250 °C for 16 min. Helium was used as a carrier gas, at a flow rate of 1.0 ml/min and mass spectra were recorded in the scan mode. The ionization voltage was 70 eV. The split ratio was 1:20. The ion source temperature was 175 °C, Interface temperature was 250 °C and 280°C. The constituents of essential oil were identified based on their retention time (R_t_) with respect to the reference. The scan range was 40-700 m/z. The identification of compounds was based on matching unknown peaks with MS-data bank (NIST 2.0 electronic Library).

### *In vitro* antioxidant activity of CEO

To analyze the antioxidant potential of CEO, different method such as DPPH, FRAP and ABTS method was employed. Various concentrations of oils (5-80 µg/ml) were prepared for each antioxidant assay, while Ascorbic acid was used as standard antioxidant compound for all assays. Antioxidant capacity was expressed in terms of IC_50_ (Half maximal inhibitory concentration), lower the value of IC_50_, higher the antioxidant capacity.

### DPPH radical scavenging activity

DPPH radical scavenging activity of CEO was measured by the method described by Barros *et al*. [18] and Rolta *et al* [19]. The capability of scavenging DPPH radical was calculated using the following equation:

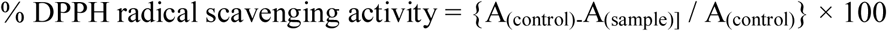

where A _(control) -_ Absorbance of control and A _(sample)_ - absorbance of the test/standard.

### FRAP activity

FRAP activity was calculated according to the method described by Benzie and Strain [20]. The antioxidant capacity of CEO and ascorbic acid was calculated from the linear calibration curve of FeSO_4_ (10 to 80 μM) and expressed as μM FeSO_4_ equivalents.

### ABTS scavenging assay

ABTS scavenging activity of CEO was calculated according to the method described by Re *et al*. [21]. Percentage ABTS scavenging activity was calculated as-

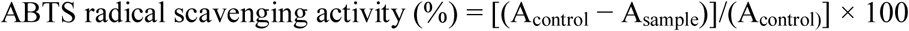

Where, A_control_ is the absorbance of ABTS radical + methanol; A_sample_ is the absorbance of ABTS radical + sample /standard.

### *In vitro* anti-inflammatory activity of CEO

CEO was investigated for its inflammatory activity using denaturation of egg albumin method as per the reported method of Chandra *et al*. [22]; Gogoi *et al*. [23]. In this method, 200 µl of egg albumin (from fresh hen’s egg), 2.8 ml of phosphate buffered saline (PBS, pH 6.4) and 2 ml of varying concentrations of the essential oil of *C. citratus* (50-400 μg/ml) was added. The reaction mixtures were then incubated at 37±2 °C in an incubator for 15 min and then heated at 70°C for 5 min in a hot water bath. After cooling down, the absorbance was measured at 660 nm against blank. A similar volume of distilled water served as control. Diclofenac sodium in the final concentration of 50-400 μg/ml was used as reference drug. The %age inhibition of protein denaturation was calculated from the following formula:

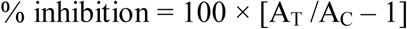

where, A_T_ = absorbance of test sample, A_C_ = absorbance of control. The experiment was done in triplicate and the average was taken.

### *In silico* prediction of bioactivity and molecular docking studies

Bioactivity potential of the major chemical constituent presents in the CEO was predicted with the help of PASS (Prediction of activity spectra for substances) prediction [24]. Molecular docking studies of the selected phytocompounds were also performed on Autodock vina [25] using enzyme Human peroxiredoxin 5 (PDB ID: 1HD2) [26] and anti-inflammatory protein, Human Cyclooxygenase-2 (5IKQ) [27] in order to know the binding affinity and various ligands. Both the proteins were retrieved from protein data bank (https://www.rcsb.org/) in pdb format. Ascorbic acid and Tocopherol were used as standard drug for antioxidant protein target, while Arachidonic acid and Diclofenac were used as standard drug for inflammatory protein target. Lipinski’s rule [28], [29] (rule of five, RO5) was to evaluate the drug-likeness property. The admet SAR [30], [31] and Protox-II server [32] were used to predict ADME and toxicity respectively. Detailed visualization and comparison of the docked sites of target proteins and ligands were done by Chimera [33] and LigPlot [34].

### Statistical analysis

The data was expressed as mean ± standard deviation, calculated using Microsoft Office Excel. The experiments were performed in triplicate and their average mean was calculated. IC_50_ values were calculated from the linear regression method.

## Results

### Percentage yield and chemical composition of CEO

The percentage yield of essential oil from fresh leaves was 0.16 ± 0.086%. GC-MS analysis of the essential oil of *C. citratus* showed the presence of 48 phytocompounds. These phytocompounds were identified by comparing the mass spectra of the constituents with the NIST mass spectral library (https://chemdata.nist.gov/) and are summarized in table 1. The mass spectra of all the phytochemicals identified in the essential oil of *C. citratus* are presented in Fig.1. Among all phytocompounds, 8-methyl-3,7-Nonadien-2-one (E) (27.28%), α-Pinene (15.60%), limonene (4.88%), citral (4.87%), limonene oxide (4.27%) and Epoxy-α-terpenyl acetate (4.03%) were major constituents, contribute 66.66% of total volatile constituents. The structures of major phytocompounds were drawn through ChemDraw software and were shown in Fig. 2.

**Table 1:**
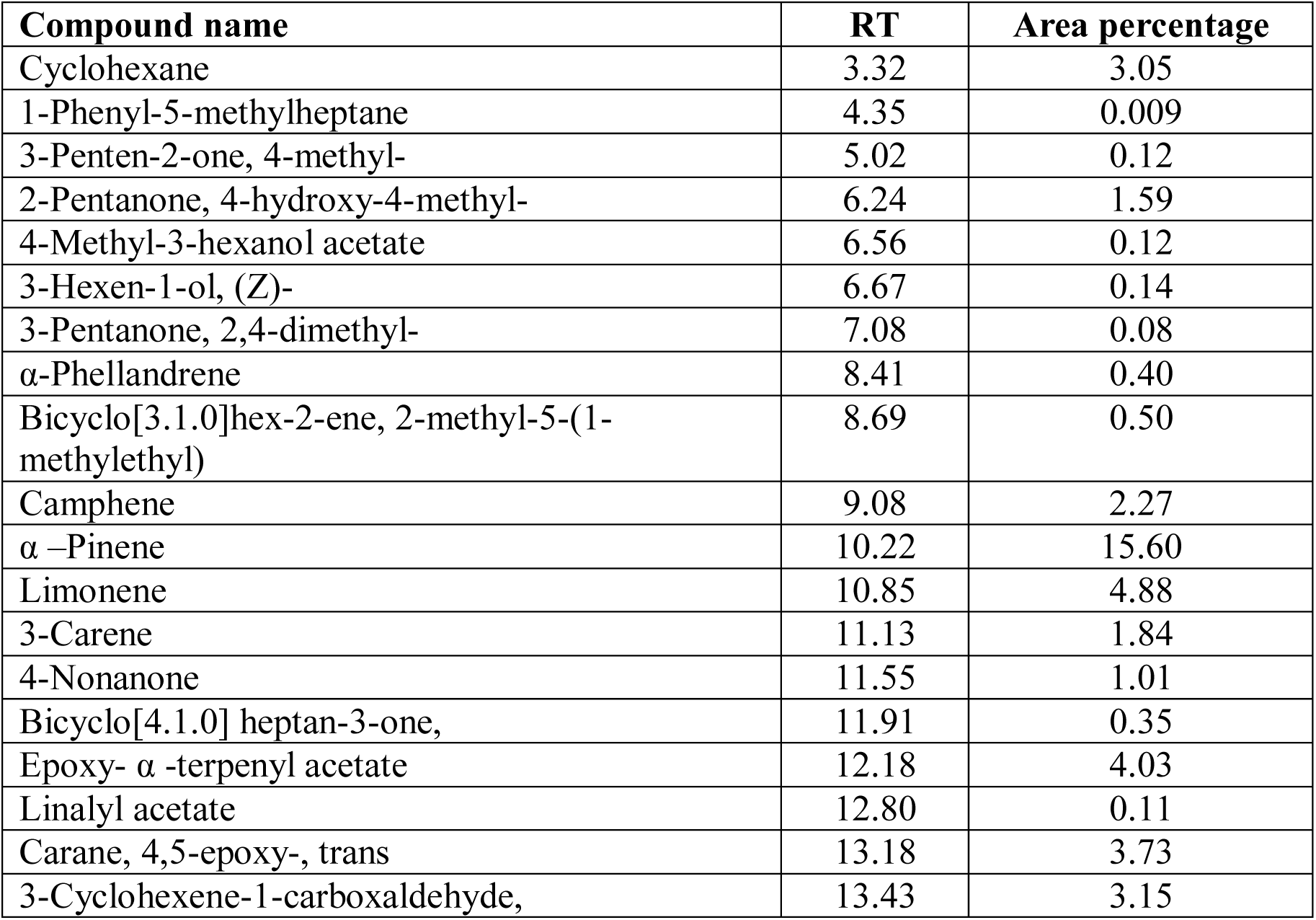
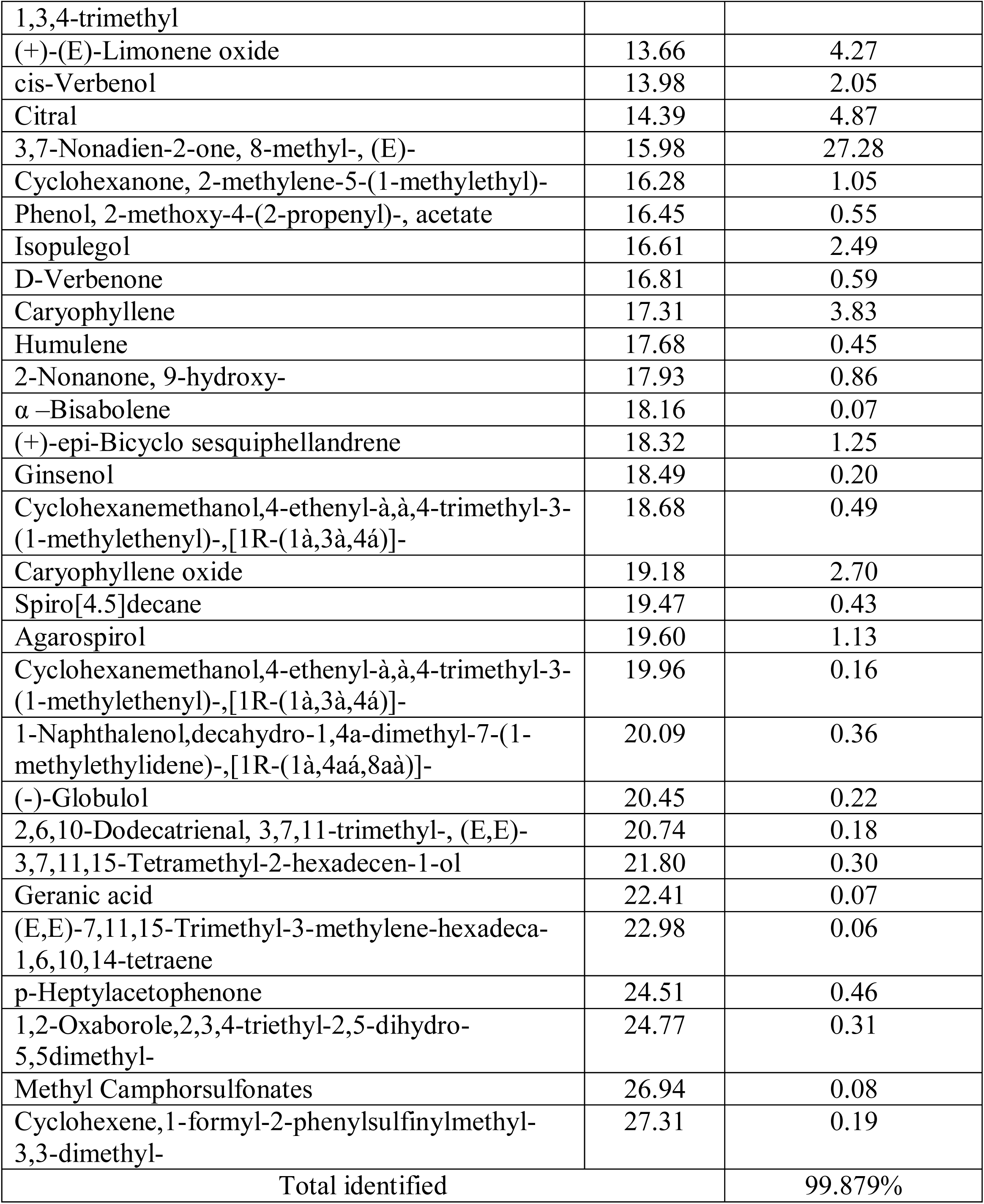
Phytocompounds identified in CEO using GC-MS analysis.

**Fig. 1:**
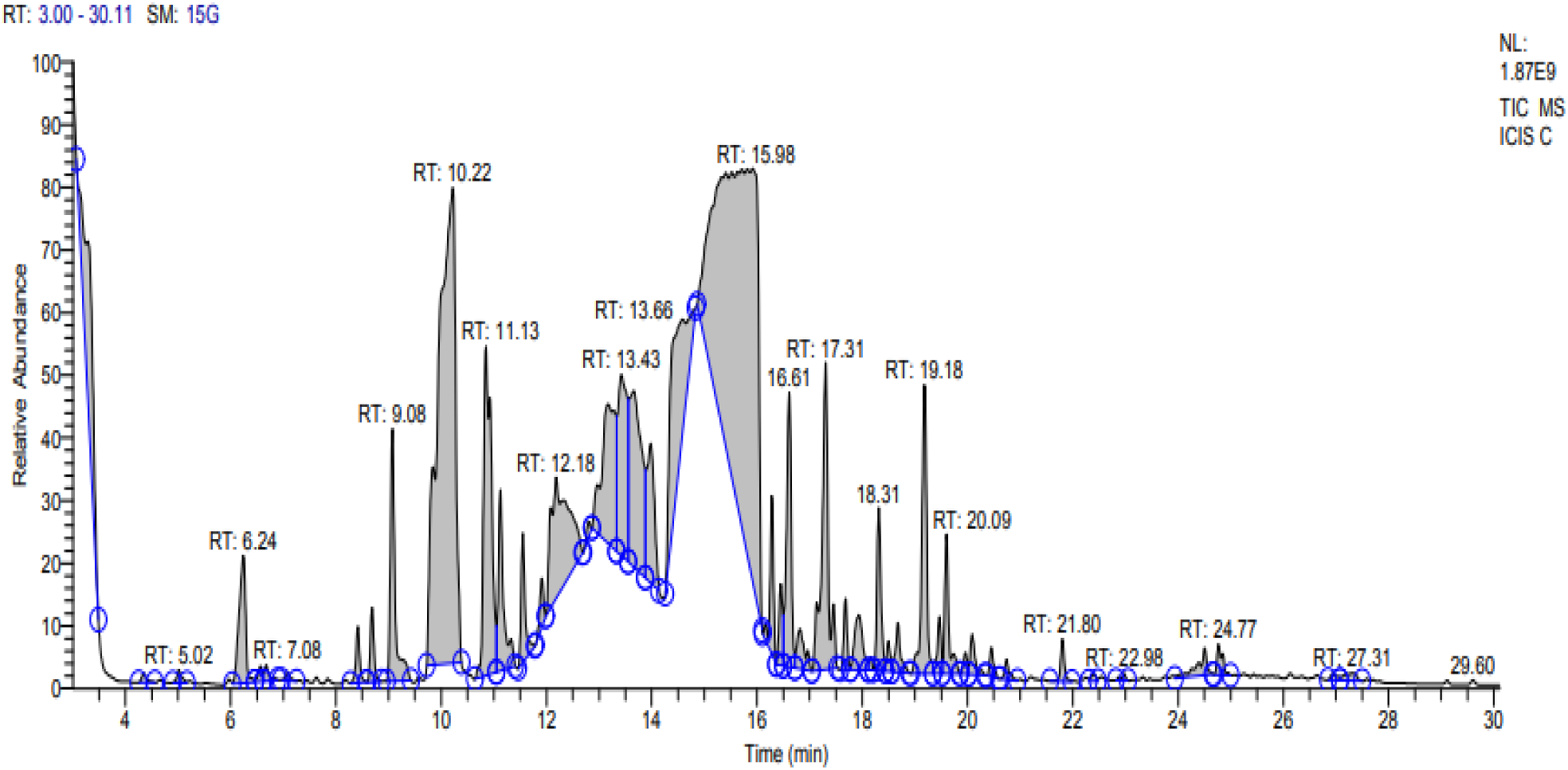
Chemical composition of CEO using GC-MS analysis.

**Fig. 2:**
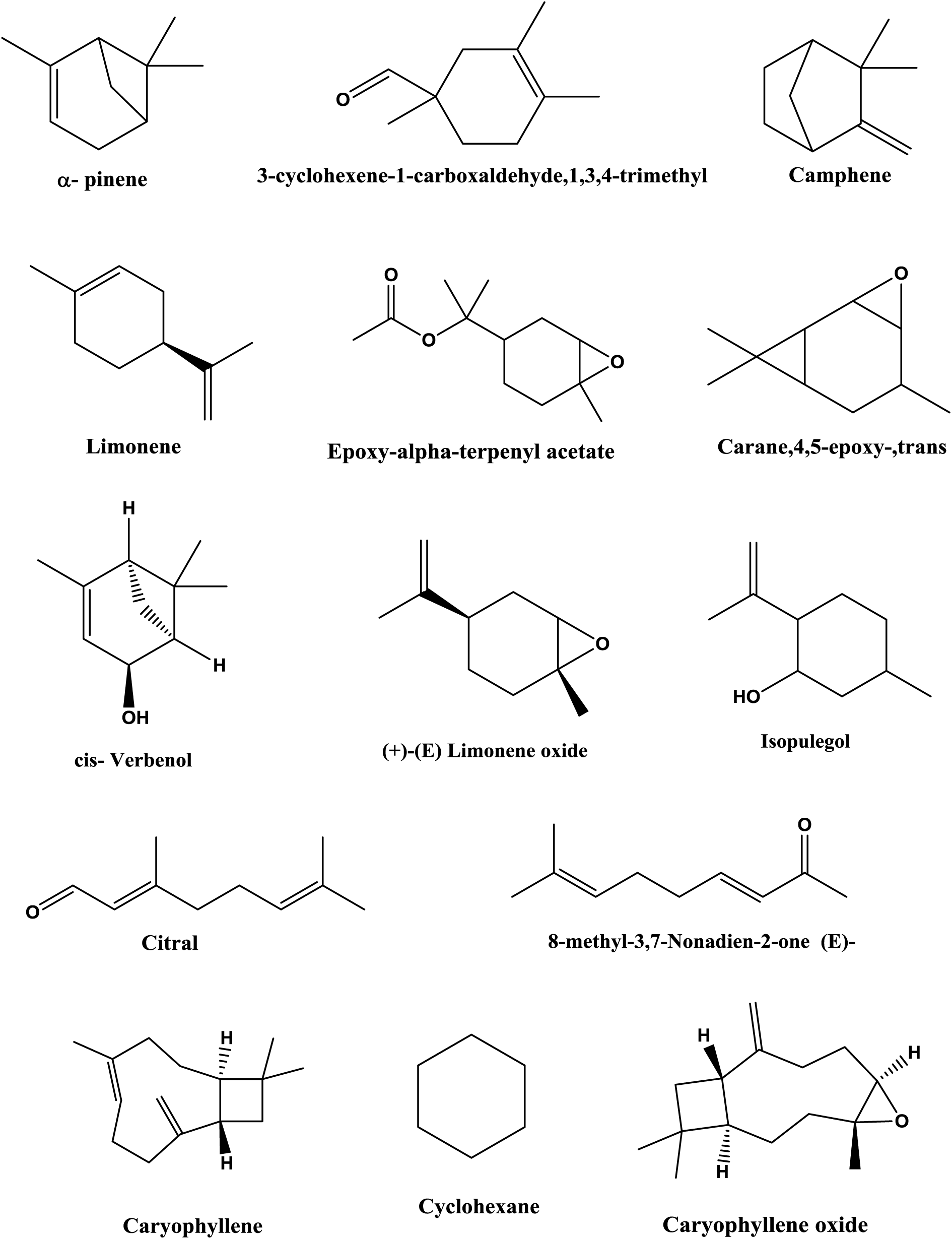
Structure of important phytocompounds identified in GC-MS analysis of CEO.

### Analysis of antioxidant activity of CEO

Essential oil of *C. citratus* (CEO) leaves followed dose-dependent pattern for all antioxidant assays (DPPH, ABTS and FRAP) as shown in Fig. 3A, B, C. IC_50_ of CEO was found to be 91.0 ± 9.25 µg ml^-1^, 350.957 ± 8.92 µM, and 370.2 ± 11.81 µg ml^-1^, with DPPH, FRAP and ABTS assay, respectively indicating strong DPPH radical scavenging activity as compared to FRAP and ABTS activity. Ascorbic acid showed IC_50_ value of 33.38 ± 2.29 µg ml^-1^, 157.26 ± µM and 170.41 ± 7.91 µg ml^-1^, respectively with DPPH, FRAP and ABTS method (Table-2).

**Fig. 3:**
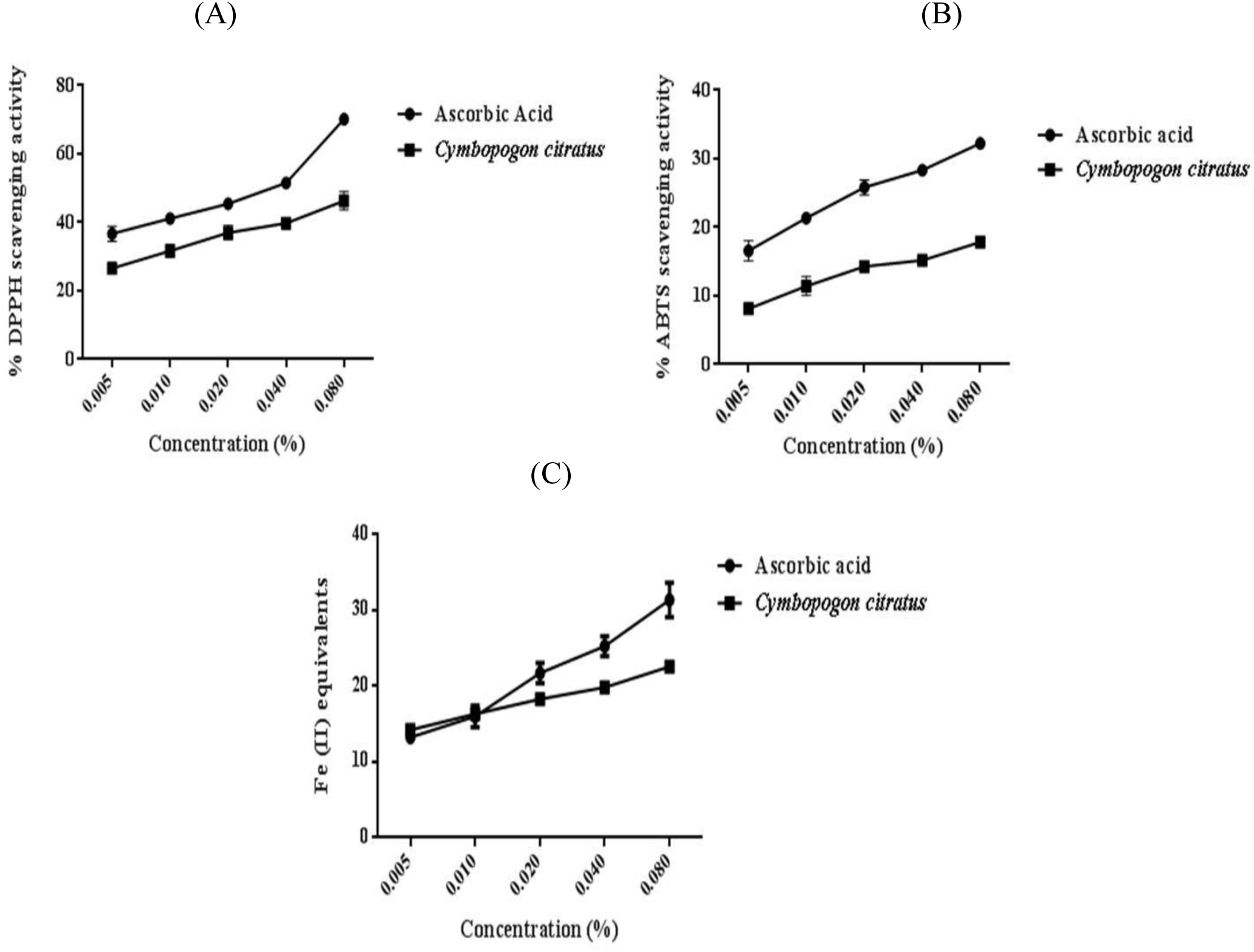
Concentration dependent antioxidant activity of CEO using DPPH radical scavenging assay (A), ABTS scavenging activity (B) and FRAP assay (C).

**Table 2:** Half maximal inhibitory concentration (IC_50_) of CEO. Ascorbic acid was used as positive control. Values are expressed as mean ± S.D. of three independent experiments.

| Antioxidant assays | IC<sub>50</sub> value |  |
| --- | --- | --- |
|  | Ascorbic acid | CEO |
| **DPPH\*** | 33.38±2.29 | 91.0±9.25 |
| **FRAP\*\*** | 157.26±19.09 | 350.957±8.92 |
| **ABTS\*** | 170.41±7.91 | 370.1985±11.81 |

\*- μg/ml; \*\*- μM Fe (II) equivalents

### Anti-inflammatory activity of CEO

*In vitro* anti-inflammatory activity of CEO was determined by denaturation of egg albumin using different concentrations (50-400 μg ml^2^) of CEO and diclofenac sodium and it showed concentration-dependent pattern of denaturation. It was observed that CEO was found to exhibit strong inflammatory activity (IC_50_-397.11± 1.45µg/ml) as compared to diclofenac (IC_50_-682.98±7.47 µg/ml) (Fig. 4).

**Fig. 4:**
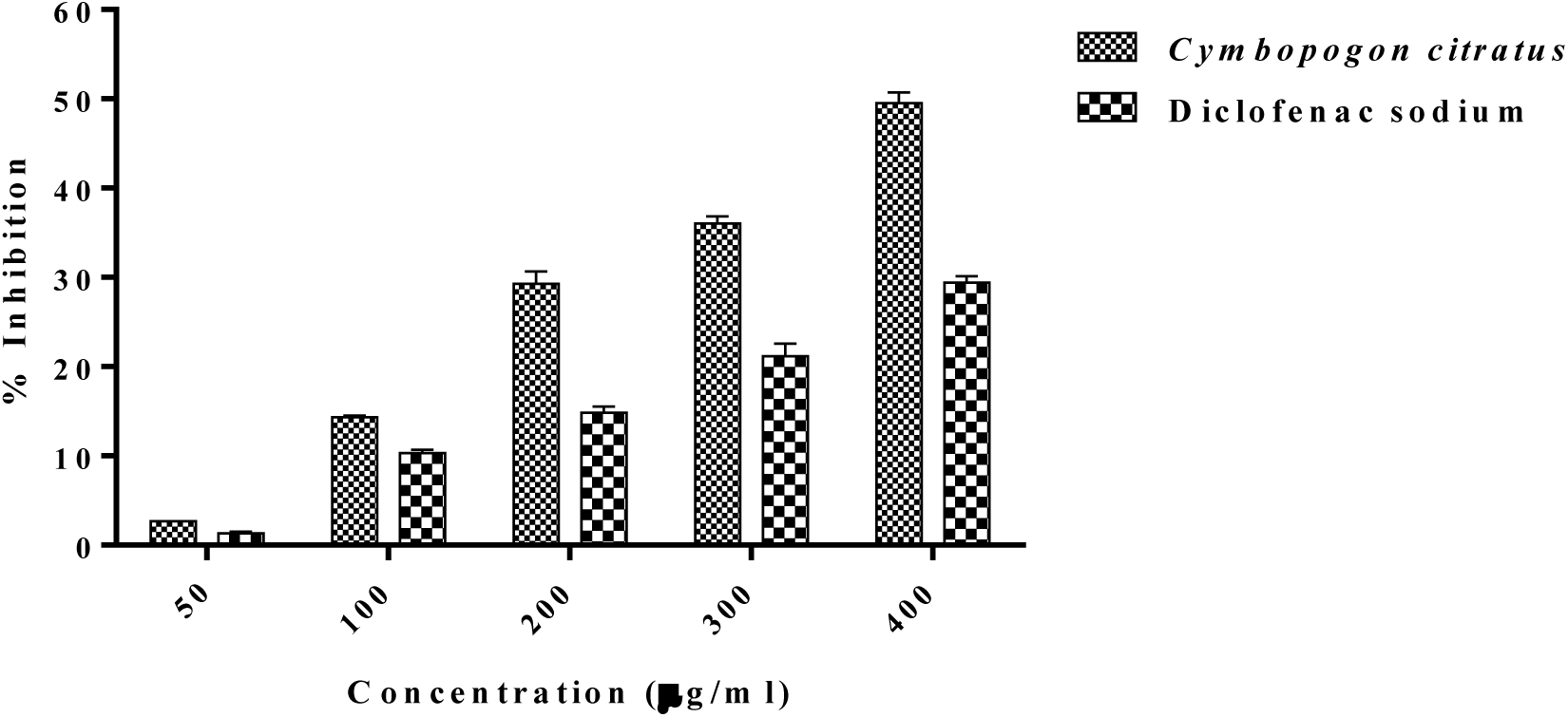
*In vitro* anti-inflammatory activity of CEO using egg albumin denaturation. Diclofenac sodium was used as control.

### Receptor-ligands interactions

Cyclooxygenase plays a major role in inflammation and is responsible for conversion of arachidonic acid to prostaglandins. It exists in two isoforms-cyclo oxygenase-1 which is constitutive and cyclo oxygenase-2 (COX-2) which is induced by cytokines [35]. Selective inhibitors of COX-2 also increase the risk of vascular events [36]. Human PRDX5 antioxidant enzyme permits the reduction of hydrogen peroxide and alkyl peroxide, with the help of thiol -containing donor molecules [37, 38]. The results of docking interaction between selected phytocompounds and targeted receptor proteins, Human peroxiredoxin 5(1HD2) and Human Cyclooxygenase-2 (5IKQ) were shown in Table 3. It was found that β-Caryophyllene showed best interaction with 1HD2 with docking score (−7.9 kcal mol^-1^) followed by caryophyllene oxide (−7.1 kcal mol^-1^) as compared to α-Tocopherol (−7.3 kcal mol^-1^) and ascorbic acid (−4.9 kcal mol^-1^). Similarly, caryophyllene oxide (−10.3 kcal/mol) and β-Caryophyllene (−10.2 kcal mol^-1^) showed highest binding energy as compared to that of diclofenac (−8.7 kcal mol^-1^) and arachidonic acid -7.0 kcal mol^-1^) (Table 3). The interacting amino acids showing H-bonding and hydrophobic interaction between phytocompounds and receptors were shown in table 3. Interactions of both the receptors with caryophyllene oxide and β-Caryophyllene were shown in Fig. 5 and 6.

**Table 3:** Binding energy calculated through Autodock vina and interactions of phytocompounds and selected drugs with target protein receptors. Diclofenac and Arachidonic acid were used as standard control with 5IKQ, while Ascorbic acid and α-Tocopherol were used as standard control with 1HD2 receptor.

| Compounds/Drugs | 1HD2 |  |  | 5IKQ |  |  |
| --- | --- | --- | --- | --- | --- | --- |
|  | Affinity (kcal/mol) | H-bonding | Non-covalent interactions | Affinity (kcal/mol) | H-bonding | Non-covalent interactions |
| 3,7-Nonadien-2-one, 8-methyl-, (E)- | -4.5 | Ser 115(A) | Asp 113(A), Leu 116(A), Leu 112(A), Thr 147(A) | -6.4 | - | Leu 352(A), Ser 353(A), Trp 387(A), Phe 518(A), Val 523(A), Gly 526(A), Leu 384(A), Tyr 385(A), Met 522(A) |
| α-pinene | -5.0 | - | Pro 40(A), Pro 45(A), Leu 116(A), Ile 119(A), Phe 120(A), Thr 44(A), Thr 147(A) | -6.7 | - | Gln 203(A), Tyr 385(A), Trp 387(A), Leu 390(A), Ala 199(A), Ala 202(A), Leu 391(A) |
| Limonene | -5.1 | - | Phe 15(A), Pro 19(A), Ala 78(A), Pro 100(A) | -7.8 | - | Phe 209(A), Ser 530(A), Leu 534(A), Phe 205(A), Phe 381(A), Gly 533(A), Gly 227(A), Val 228(A), Asn 375(A), Ile 377(A) |
| Citral | -4.6 | Thr 147(A) | Phe 43(A), Val 80(A), Leu 149(A), Gly 46(A), Ala 42(A), Asn 76(A), Glu 83(A) | -6.3 | Thr 206(A) | Trp 387(A), Gln 203(A), Ala 202(A), Ala 199(A), Tyr 385(A), Leu 390(A), His 207(A), Leu 391(A) |
| Epoxy- $\alpha$ -terpenyl acetate | -5.5 | - | Pro 40(A), Thr 147(A), Leu 116(A), Phe 120(A), Thr 44(A), Ile 119(A), Arg 127(A) | -7.9 | - | Val 523(A), Leu 352(A), Gly 526(A), Ser 353(A), Phe 518(A), Met 522(A), Ala 527(A), Trp 387(A) |
| β-Caryophyllene | -7.9 | - | Pro 19(A), Asp 77(A), Ala 78(A), Pro 100(A), Phe 15(A) | -10.2 | - | Val 523(A), Phe 381(A), Leu 384(A), Tyr 385(A), Trp 387(A), Met 522(A), Ala 527(A), Ser 530(A), Gly 526(A), Phe 518(A) |
| Carane, 4,5-epoxy-, trans | -5.5 | - | Phe 15(A), Pro 19(A), Asp 77(A), Ala 78(A), Pro 100(A) | -7.7 | Trp 387(A) | Leu 391(A), Ala 199(A), Ala 202(A), Leu 390(A), Tyr 385(A) |
| 3-Cyclohexene-1-carboxaldehyde, 1,3,4-trimethyl | -5.7 | Ala 78(A) | Pro 19(A), Asp 77(A), Pro 100(A), Phe 15(A) | -7.2 | - | Gln 203(A), Tyr 385(A), Ala 202(A), Ala 199(A), Thr 206(A), Trp 387(A), Leu 390(A) |
| Cyclohexane | -4.1 | - | Pro 19(A), Phe 15(A), Asp 77(A), Ala 78(A), Pro 100(A) | -5.3 | - | Ser 530(A), Phe 205(A), Phe 209(A), Phe 381(A), Gly 533(A), Leu 534(A), Val 228(A) |
| Caryophyllene oxide | -7.1 | - | Pro 40(A), Thr 147(A), Thr 44(A), Phe 120(A), Pro 45(A), Leu 116(A), Ile 119(A) | -7.2 | - | Tyr 385(A), Phe 518(A), Val 523(A), Val 349(A), Phe 381(A), Leu 384(A), Trp 387(A), Ser 530(A), Gly 526(A), Ala 527(A), Met 522(A) |
| Isopulegol | -5.1 | - | Phe 15(A), Pro 19(A), Ala 78(A),<br>Pro 100(A), Asp 77(A) | -7.2 | Tyr 385(A),<br>Trp 387(A) | Ala 202(A), Gln 203(A), Ala 199(A), Thr<br>206(A), Leu 391(A) |
| Camphene | -5.2 | - | Pro 19(A), Asp 77(A), Ala 78(A),<br>Pro 100(A), Phe 15(A) | -7.0 | - | Phe 381(A), Leu 384(A), Tyr 385(A), Trp<br>387(A), Met 522(A), Ser 530(A), Gly 526(A),<br>Phe 518(A), Val 523(A), Leu 352(A) |
| Cis-verbenol | -5.0 | - | Phe 15(A), Pro 19(A), Asp 77(A),<br>Ala 78(A), Pro 100(A) | -7.2 | - | Phe 518(A), Met 522(A), Leu 352(A), Leu<br>384(A), Tyr 385(A), Trp 387(A), Val 523(A),<br>Ser 530(A), Gly 526(A) |
| Diclofenac | - | - | - | -8.7 | - | Tyr 385(A), Trp 387(A), His 388(A), Leu<br>391(A), Thr 206(A), His 207(A), Ala 202(A),<br>Gln 203(A), Leu 390(A) |
| Arachidonic acid | - | - | - | -7.0 | Tyr 355(A) | Tyr 385(A), Arg 120(A), Val 349(A), Leu<br>352(A), Ser 353(A), Trp 387(A), Phe 518(A),<br>Ala 527(A), Ser 530(A), Met 522(A), Gly<br>526(A), Val 523(A) |
| Ascorbic acid | -4.9 | Asn 122(A),<br>Asp 77(A),<br>Arg 124(A),<br>Asn 76(A) | Pro 100 (A), Val 75(A) | - | - | - |
| α-Tocopherol | -7.2 | Arg 127(A) | Asp 113(A), Thr 147(A), Leu<br>116(A), Ser 115(A), Leu 112(A),<br>Pro 40(A), Thr 44(A), Gly 46(A),<br>Cys 47(A), Phe 120(A), Asp<br>145(A) | - | - | - |

**Fig. 5:**
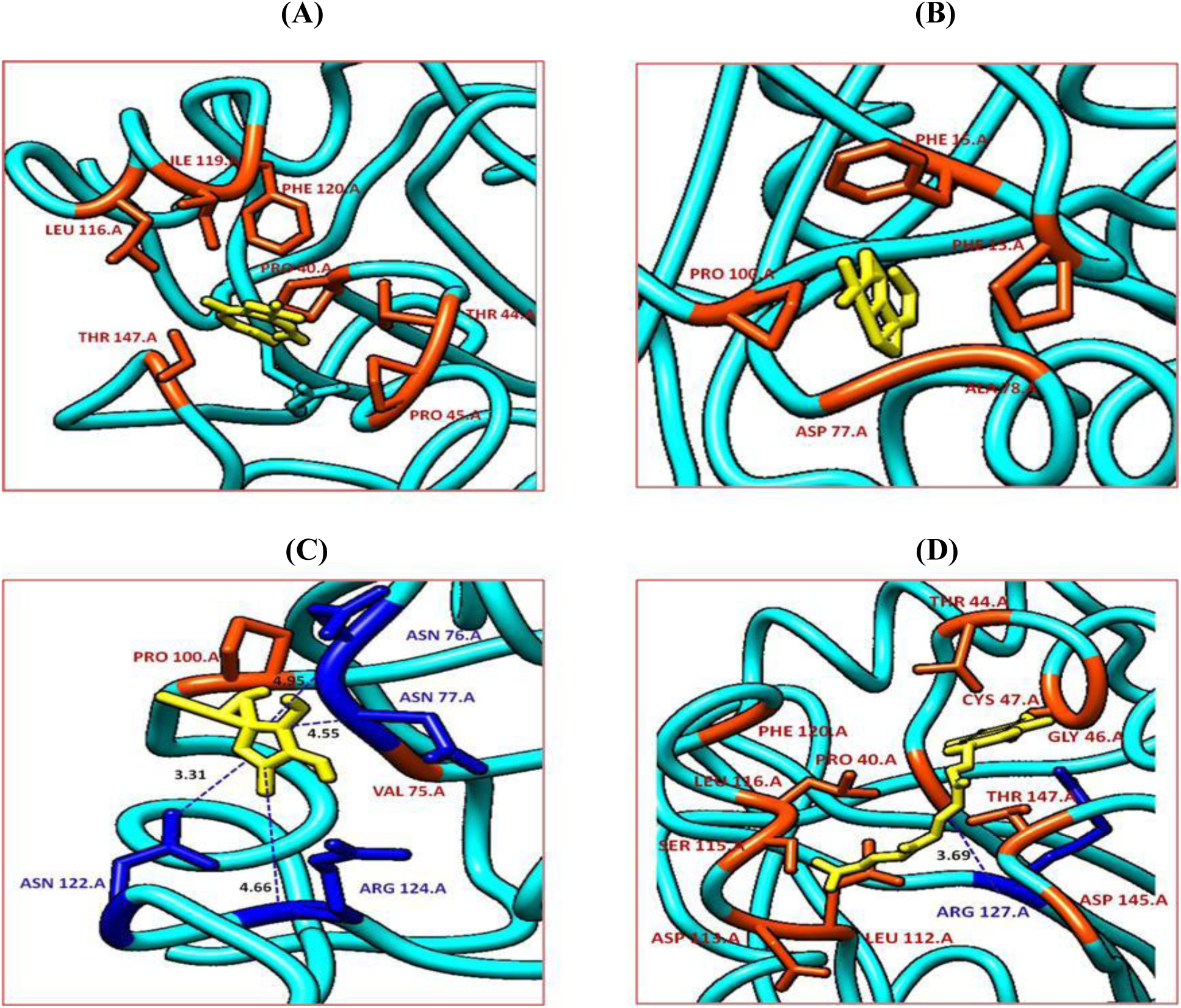
Interactions of Human Peroxiredoxin receptor with caryophyllene oxide (A), β-caryophyllene (B), ascorbic acid (C) and α-Tocopherol (D). Amino acids with hydrophobic interactions were shown in orange red color, whereas, amino acids showing H-bonding with receptor were shown in blue color.

**Fig. 6:**
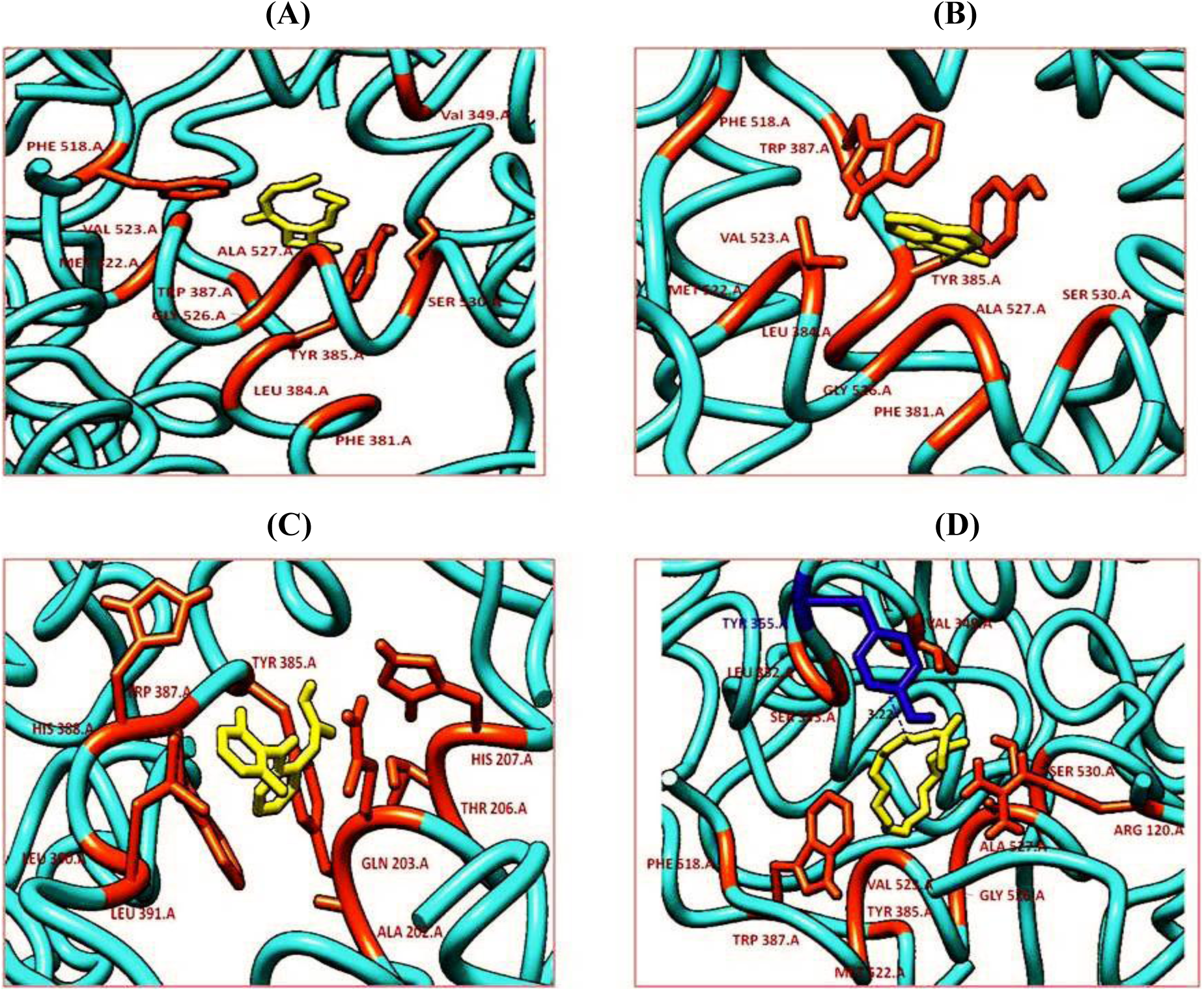
Interactions of Human Cyclooxygenase-2 receptor with caryophyllene oxide (A), β-caryophyllene (B), Diclofenac (C) and Arachidonic acid (D). Amino acids with hydrophobic interactions were shown in orange red color, whereas, amino acids showing H-bonding with receptor were shown in blue color.

LigPlot structures of all the selected phytocompounds with both the target receptors were shown in supplementary data (Supplementary Fig. 1 and Supplementary Fig. 2).

### Drug likeness prediction of selected phytocompounds of CEO

The drug likeness filters help in the early preclinical development by avoiding costly late step preclinical and clinical failure. The drug likeness properties of phytocompounds showing good interactions (α-Pinene, Limonene, (+)-(E)-Limonene oxide, Isopulegol, Caryophyllene oxide) were analyzed based on the Lipinski rule of 5. It was found that all the selected phytocompounds followed Lipinski’s rule of five (Table 4).

**Table 4:** *In silico* PASS prediction for anti-inflammatory and antioxidant activity of selected phytocompounds of CEO.

| Phytocompounds/drugs | Anti-inflammatory |  | Antioxidant |  |
| --- | --- | --- | --- | --- |
| | $P_a$ | $P_i$ | $P_a$ | $P_i$ |
| α-Pinene | 0.490 | 0.060 | - | - |
| Limonene | 0.610 | 0.029 | 0.157 | 0.094 |
| Limonene oxide | 0.654 | 0.022 | 0.153 | 0.099 |
| β-Caryophyllene | 0.745 | 0.011 | 0.174 | 0.074 |
| Caryophyllene oxide | 0.759 | 0.009 | 0.144 | 0.110 |
| Isopulegol | 0.690 | 0.017 | 0.184 | 0.066 |
| Diclofenac | 0.791 | 0.007 | - | - |
| Arachidonic acid | 0.730 | 0.012 | - | - |
| α-Tocopherol | - | - | 0.967 | 0.002 |
| Ascorbic acid | - | - | 0.928 | 0.003 |

CEO= Essential oil of *C. citratus* leaves; $P_a$ = Probable activity; $P_i$ = probable inactivity; PASS = Prediction of Activity Spectra for Substances

**Table 2:**
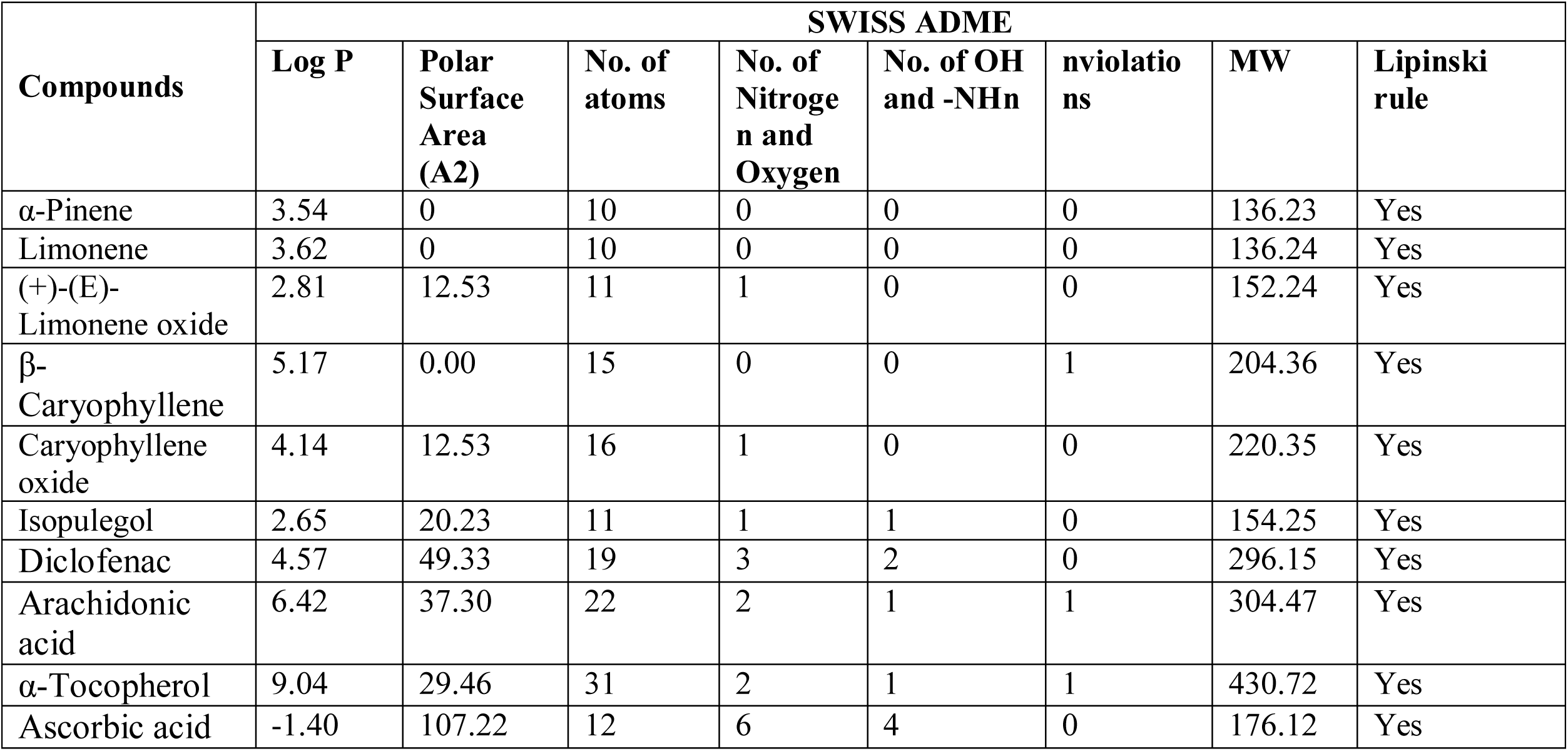
Drug-likeness prediction of selected phytocompounds from *C. citratus*

| Compounds | SWISS ADME |  |  |  |  |  |  | Lipinski rule |
| --- | --- | --- | --- | --- | --- | --- | --- | --- |
|  | Log P | Polar Surface Area (A2) | No. of atoms | No. of Nitrogen and Oxygen | No. of OH and -NHn | nviolations | MW |  |
| α-Pinene | 3.54 | 0 | 10 | 0 | 0 | 0 | 136.23 | Yes |
| Limonene | 3.62 | 0 | 10 | 0 | 0 | 0 | 136.24 | Yes |
| (+)-(E)-Limonene oxide | 2.81 | 12.53 | 11 | 1 | 0 | 0 | 152.24 | Yes |
| β-Caryophyllene | 5.17 | 0.00 | 15 | 0 | 0 | 1 | 204.36 | Yes |
| Caryophyllene oxide | 4.14 | 12.53 | 16 | 1 | 0 | 0 | 220.35 | Yes |
| Isopulegol | 2.65 | 20.23 | 11 | 1 | 1 | 0 | 154.25 | Yes |
| Diclofenac | 4.57 | 49.33 | 19 | 3 | 2 | 0 | 296.15 | Yes |
| Arachidonic acid | 6.42 | 37.30 | 22 | 2 | 1 | 1 | 304.47 | Yes |
| α-Tocopherol | 9.04 | 29.46 | 31 | 2 | 1 | 1 | 430.72 | Yes |
| Ascorbic acid | -1.40 | 107.22 | 12 | 6 | 4 | 0 | 176.12 | Yes |

### 3.3 Toxicity and ADME/T prediction of phytocompounds of *C. citratus*

The results of admetSAR analysis and toxicity prediction were shown in table 5. All of the phytochemicals showed an acceptable range of ADME/T profiles that reflect their efficiency as potent drug candidates. All the compounds showed good human intestinal solubility (HIA) and are non-carcinogenic in nature. Rat acute toxicity concentration of all the compounds was high, indicating low toxicity (Table 5). Rodent toxicity (LD_50_) values for all selected compounds were higher, indicating non-toxic nature of these compounds. Also, these compounds were non-cytotoxic and non-hepatotoxic except α-Pinene (Table 3).

**Table 3:** ADMET and Protox-II prediction of selected phytocompounds of *C. citratus* and drugs used through admetSAR and Protox-II software.

| Compounds/<br>drugs | admet SAR |  |  | Protox II |  |  |
| --- | --- | --- | --- | --- | --- | --- |
|  | Human<br>intestinal<br>absorption | Carcinogens | Rate Acute<br>Toxicity (LD<sub>50</sub>)<br>kg/mol | LD<sub>50</sub> (mg<br>kg<sup>-1</sup>) | Hepatotoxicity | Cytotoxicity |
| α-Pinene | Positive | Non-Carcinogens | 1.527 (III) | 1190<br>(class 4) | Active | Inactive |
| Limonene | Positive | Non-Carcinogens | 1.856 (III) | 4400<br>(class 5) | Inactive | Inactive |
| (+)-(E)-<br>Limonene oxide | Positive | Non-Carcinogens | 1.438 (III) | 5000<br>(class 5) | Inactive | Inactive |
| β-<br>Caryophyllene | Positive | Non-Carcinogens | 2.366 (III) | 5000 (class<br>5) | Inactive | Inactive |
| Caryophyllene<br>oxide | Positive | Non-Carcinogens | 2.236 (III) | 5000<br>(class 5) | Inactive | Inactive |
| Isopulegol | Positive | Non-Carcinogens | 2.923 (III) | 2000<br>(class 4) | Inactive | Inactive |
| Diclofenac | Positive | Non-Carcinogens | 3.029 (II) | 53 (class 3) | Active | Inactive |
| Arachidonic<br>acid | Positive | Non-Carcinogens | 1.782 (IV) | 10000<br>(class 6) | Inactive | Inactive |
| α-Tocopherol | Positive | Non-Carcinogens | 2.47 (III) | 5000 (class<br>5) | Inactive | Inactive |
| Ascorbic acid | Positive | Non-Carcinogens | 0.481 (IV) | 3367 (class<br>5) | Inactive | Inactive |

### 3.4 *In silico* PASS prediction of selected phytocompounds of CEO

The selected phytocompounds of CEO were evaluated for their anti-inflammatory and antioxidant spectra, and results of PASS prediction was shown in table 4. In the case of anti-inflammatory activity, all the compounds showed greater P_a_ than 0.5, except α-Pinene. Caryophyllene oxide and β-Caryophyllene showed highest P_a_ value of 0.759 and 0.745; while in case of antioxidant activity, all the selected phytocompounds showed P_a_ <0.5. Among all compounds, Isopulegol and β-Caryophyllene showed highest P_a_ value of 0.184 and 0.174 respectively (Table 4).

## Discussions

Traditional medicines play paramount role to cure different diseases. The plants utilized as a medicine from pre-historic time plays consequential role in primary health care. Medicinal plant contains variants of secondary metabolites which are responsible for pharmacological activities [39]. Several previous reports showed the use of plant as medicine to cure inflammation exhibiting antioxidant and anti-inflammatory activities [40]. Antioxidant compounds have been reported to obviate oxidative stress of free radicals associated with pathogenesis of a number of chronic diseases including diabetes and inflammation [41]. In present investigation, we have reported 48 phytocompounds in essential oil of *C. citrates*, out of which, 8-methyl-3,7-Nonadien-2-one (E) (27.28%) was the major constituents.

Several studies have been reported chemical composition of essential oil from *C. citrates* and found geranial, neral, and myrcene as major components [42-50]. Similar results were also reported by Hanna et al. [51] while investigating the effect of drying method on chemical composition of essential oil of lemon grass; however, there was variation in number of components, but components were same with all drying methods. Longifolene-(V4) (56.67%) and selina-6-ezn-4-ol (20.03%) was found to be major constituents in essential oil obtained from roots of *C. citratus* [52]. This variation in essential oil from *C. citratus* may be attributed to different geographical location, climate conditions, harvest period, plant age and distillation method [53-54].

Antioxidants play an important role in the prevention and promotion of health in humans against the harmful free radicals that cause many age-related diseases. Similar to our report, Farias et al. [55] also found low antioxidant capacity of essential oil from *C. citratus*. There are several studies, which showed good antioxidant potential of essential oil from *C. citratus* [43], [56-59]. In addition to essential oil, several reports have described the antioxidant potential of various extracts of *C. citratus* [60-62]. The essential oil of *C. citratus* showed strong anti-inflammatory activity as compared to standard drug, Diclofenac sodium. Anti-inflammatory activity of *C. citratus* was due to the presence of luteolin glycosides [63-64]. *In vivo* topical and oral anti-inflammatory potential of lemon grass essential oil was also reported by Boukhatem et al. [65] using carrageenan-induced paw edema test and croton oil-induced ear edema in mouse model. Costa et al. [66] also reported promising topical anti-inflammatory activity of *C. citratus* infusion, containing luteolin 7-*O*-neohesperidoside, cassiaoccidentalin B, carlinoside, cynaroside and tannins. Molecular docking study with antioxidant Human peroxiredoxin 5 (PDB ID: 1HD2) and anti-inflammatory protein, Human Cyclooxygenase-2 (PDB ID: 5IKQ) receptor showed that among all selected major phytocompounds, caryophyllene showed best interaction with 1HD2 with binding energy (−7.9 kcal/mol) which is higher than tocopherol (−7.3 kcal/mol) and ascorbic acid (−4.9 kcal/mol). Similarly, with 5IKQ, caryophyllene oxide and caryophyllene showed highest binding energy (−10.3 and -10.3 kcal/mol respectively) as compared to that of diclofenac (−8.7 kcal/mol) and arachidonic acid -7.0 kcal/mol). Both the phytocompounds also qualify ADMET features. These compounds were found to be safe by admetSAR and Protox-II software and also show highest P_a_ value with anti-inflammatory. The present study is an attempt showing anti-inflammatory and antioxidant activity of CEO through *in vitro* method and inhibitory action of phytocompounds of CEO against cyclooxygenase 2 and human peroxiredoxin proteins.

## Conclusion

*C. citratus* is one of the important herbs which play an important role in human health due to the presence of phytochemicals which are responsible for its biological activity. However, the quantification of these phytochemicals in *C. citratus* is affected by geographical and climatic conditions. In the current investigation, CEO was examined for its chemical composition, in vitro antioxidant and anti-inflammatory activity, to provide a justification of its health benefits. CEO exhibit good DPPH radical scavenging activity, while FRAP and ABTS activity of CEO was very low which may be due to combined effect of several phytocompounds. However, CEO showed significant anti-inflammatory activity. *In silico* prediction and molecular docking studies showed that Caryophyllene oxide and β-Caryophyllene contributed to antioxidant and anti-inflammatory activity of CEO. However, except α-Pinene (hepatotoxic), all the phytocompounds were found to qualify ADME/T condition and are less toxic in nature. In a nutshell, the present study opens new avenues for the plant *C. citratus* to be used as safe and less toxic alternatives to synthetic drugs used in complications arising due to oxidative stress and inflammation.

