## Supplementary Fig.1 and 2. for "*In silico* and *In vitro* evaluation of the anti-inflammatory and antioxidant potential of *Cymbopogon citratus* from North-western Himalayas"

(A) **(B)	(C)**

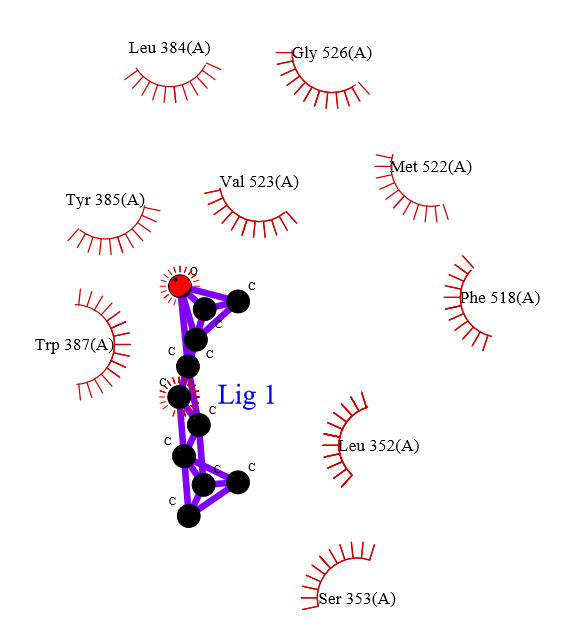

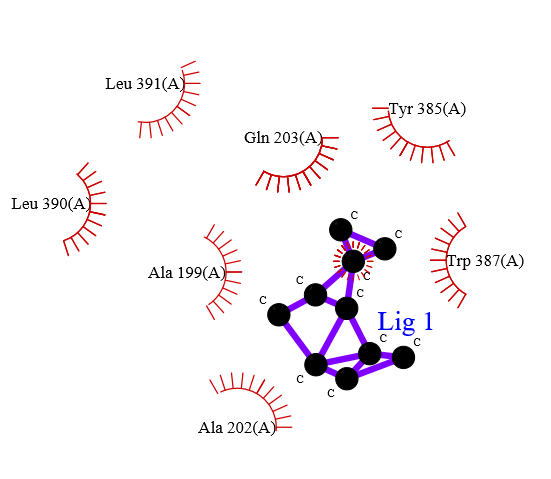

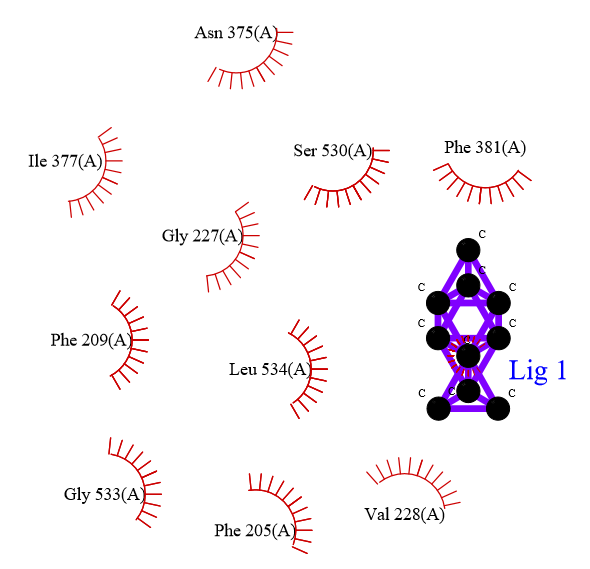

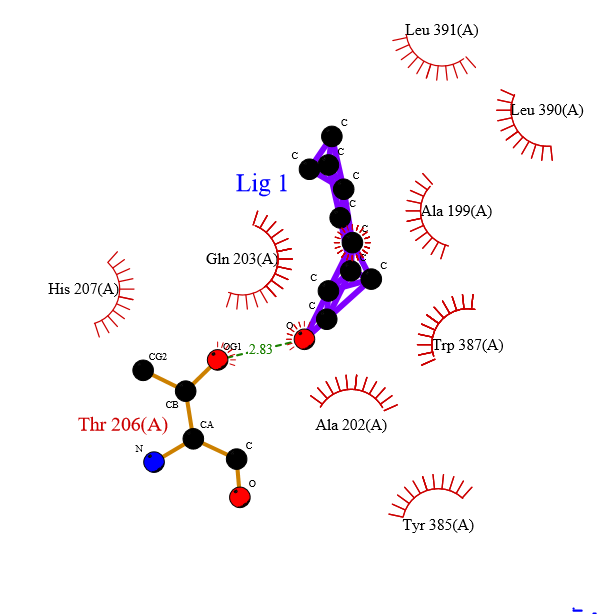

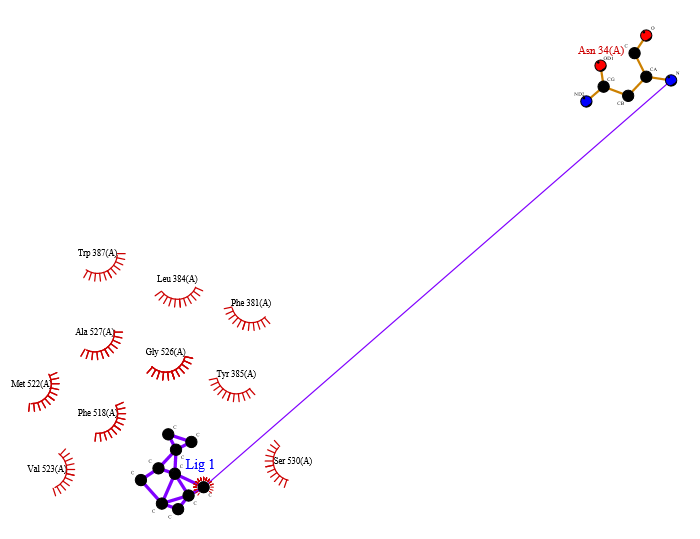

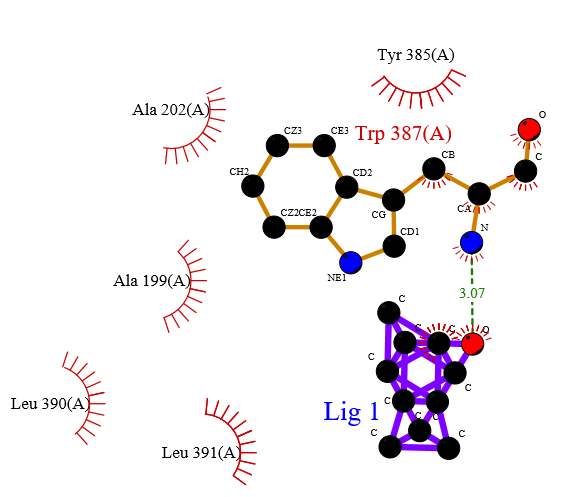

**(D)	(E)	(F)**

**(G)	(H)	(I)**

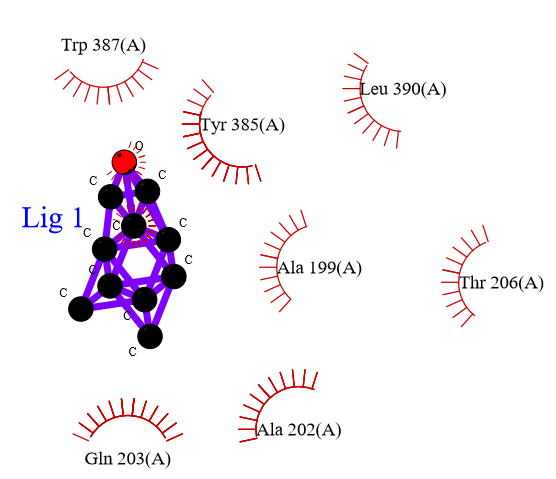

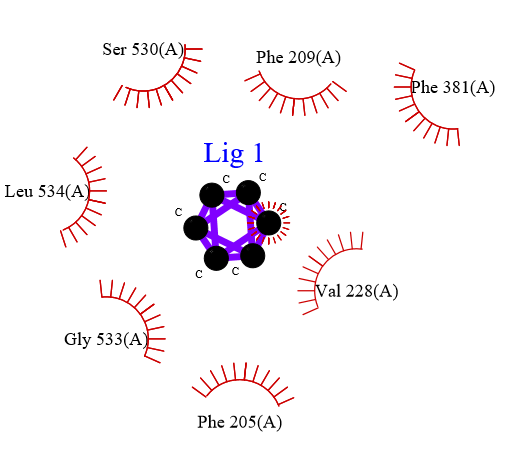

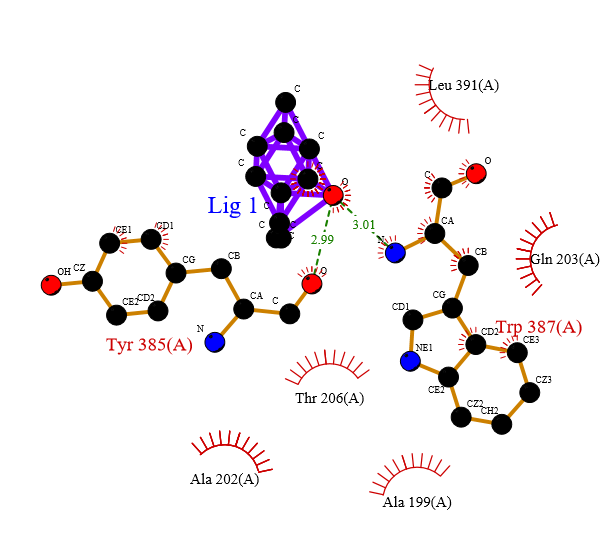

**(J)	(K)	(L)**

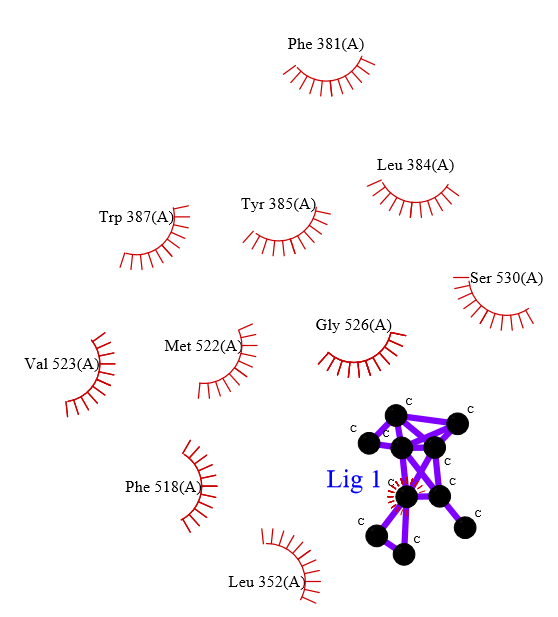

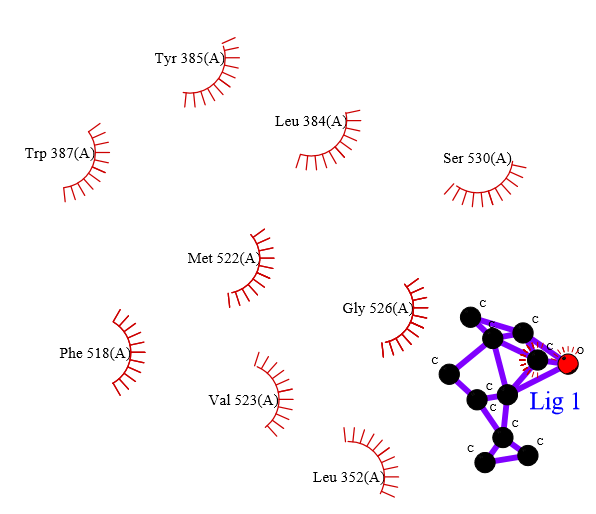

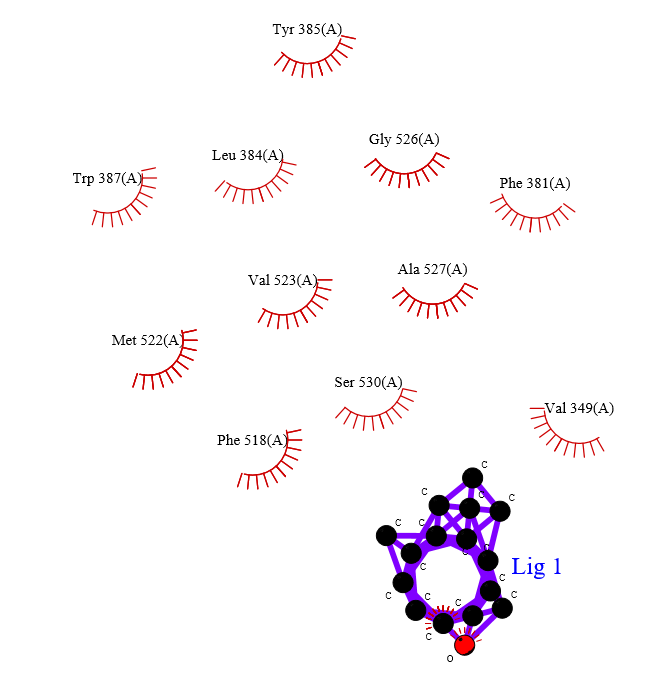

**(M)	(N)	(O)**

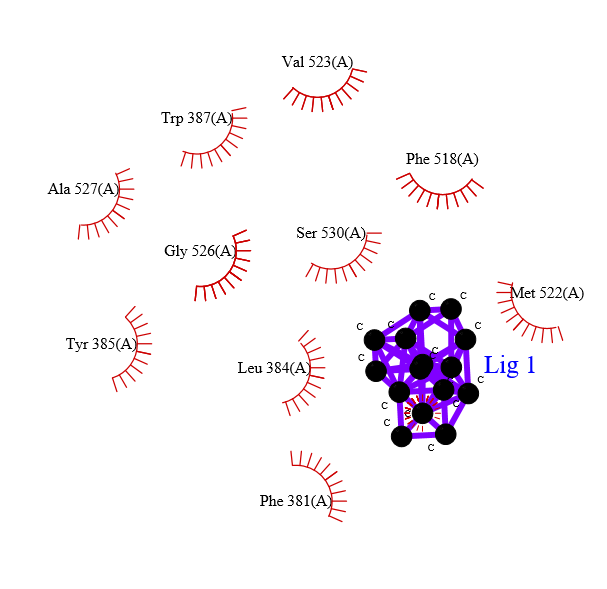

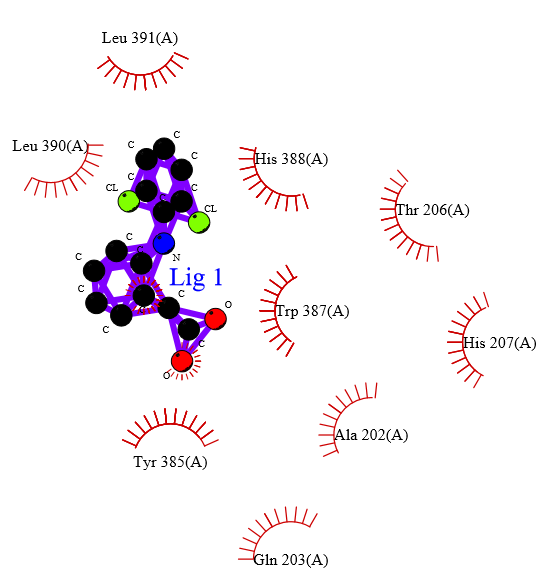

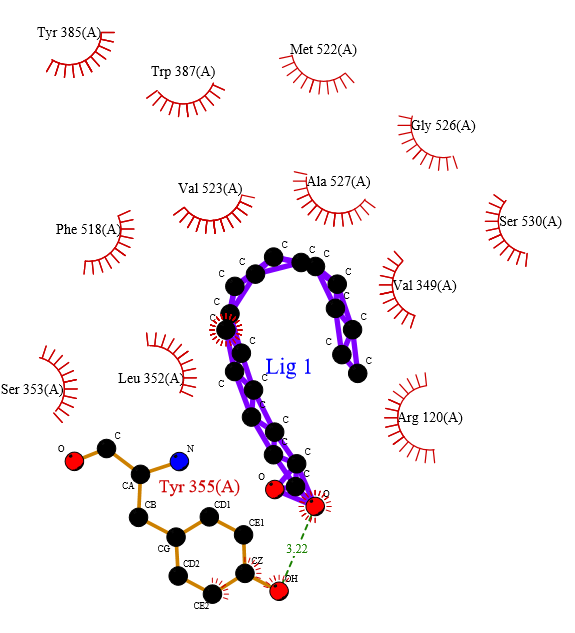

**Supplementary Fig. 1: LigPlot structure showing interactions between target protein (5IKQ) with selected phytocompounds of CEO. 3,7-Nonadien-2-one, 8-methyl-, (E)- (A), α-pinene (B), Limonene (C), Citral (D), Epoxy- α -terpenyl acetate (E), Carane (F), 4,5-epoxy-, trans (G), 3-Cyclohexene-1-carboxaldehyde,1,3,4-trimethyl (H), Cyclohexane (I), Isopulegol (J), Camphene (J), Cis-verbenol (K), Caryophyllene oxide (L), Caryophyllene (M), Diclofenac (N), Arachidonic acid (O).**

(A)	(B)	(C)

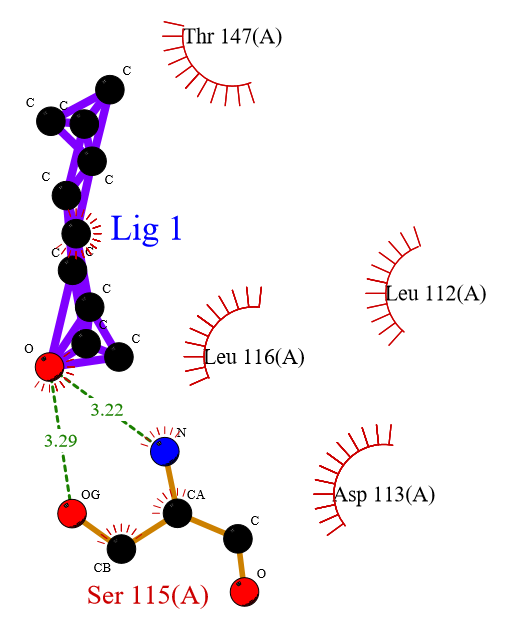

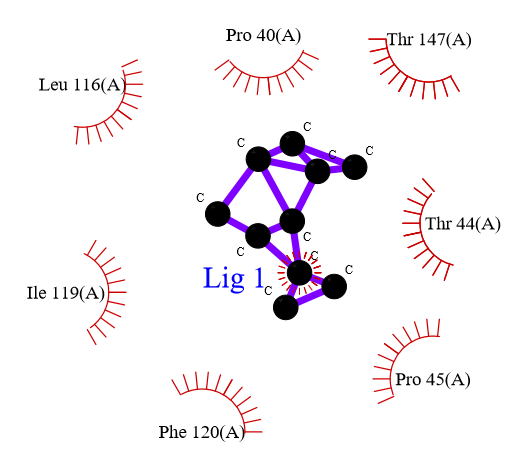

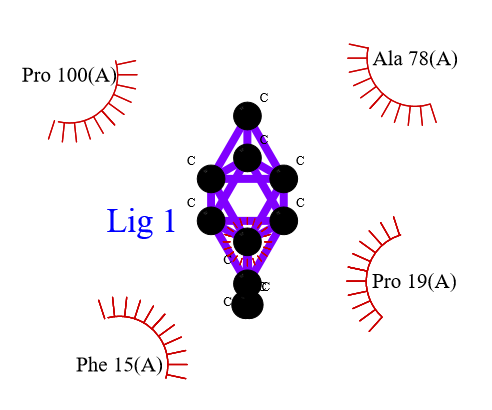

(D)	(E)	(F)

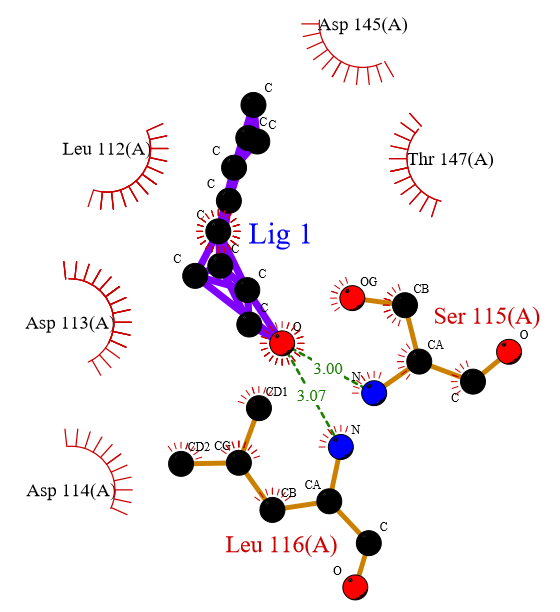

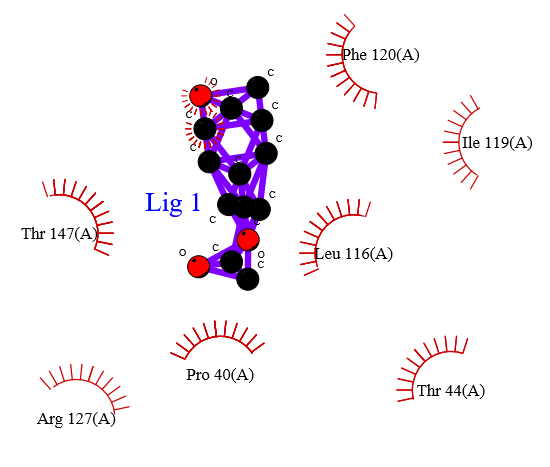

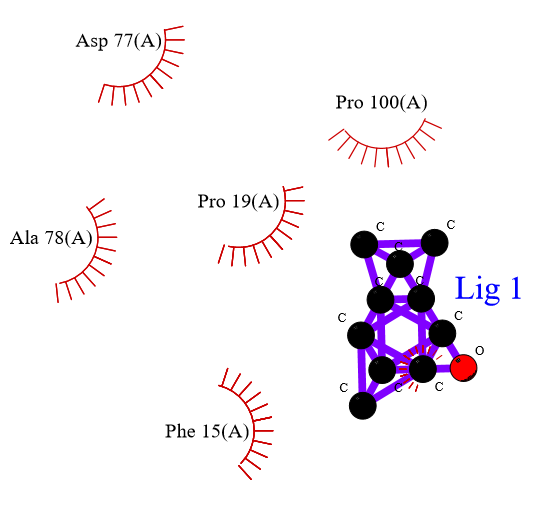

(G)	(H)	(I)

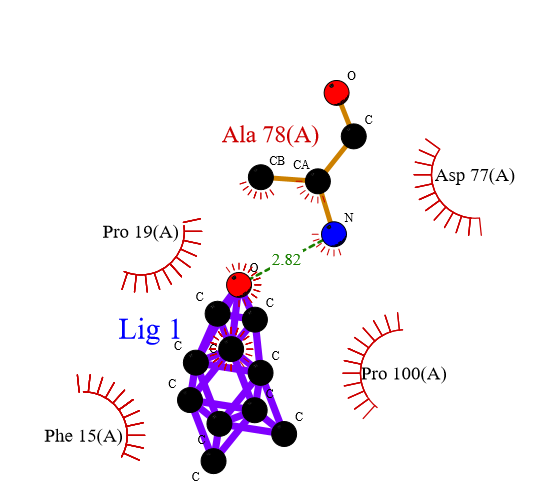

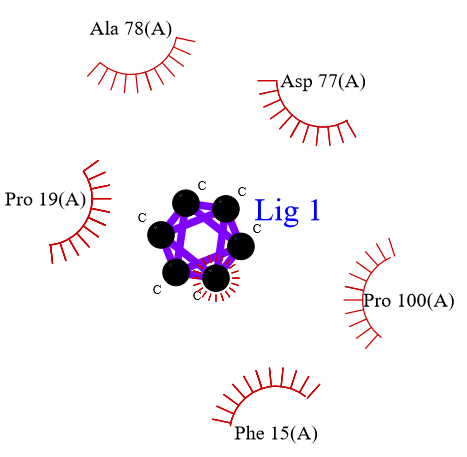

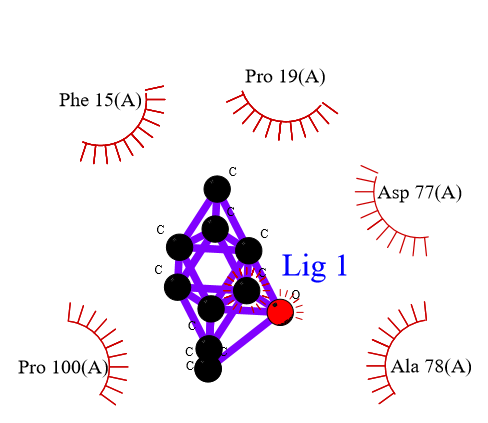

(J)	(K)	(L)

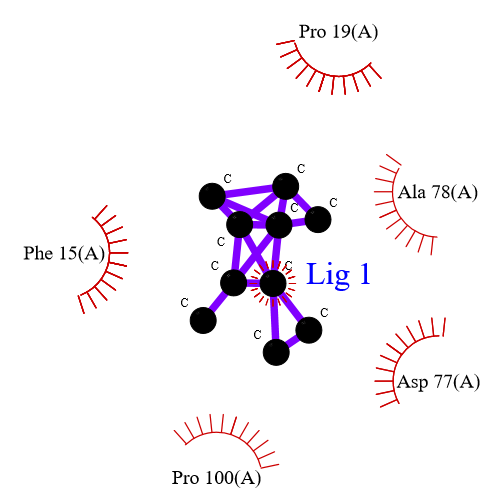

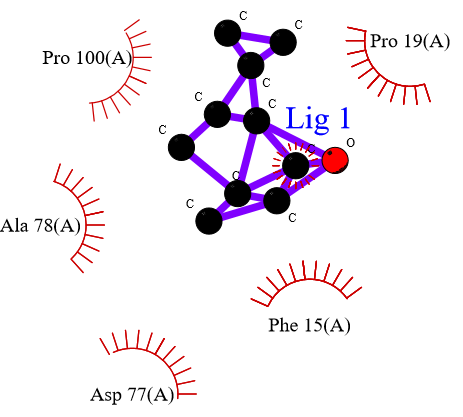

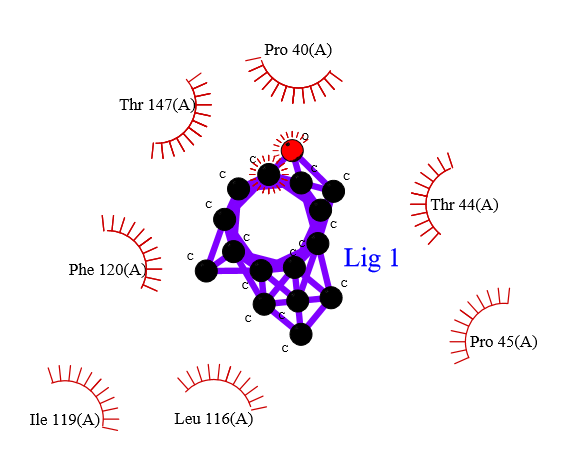

(M)	(N)	(O)

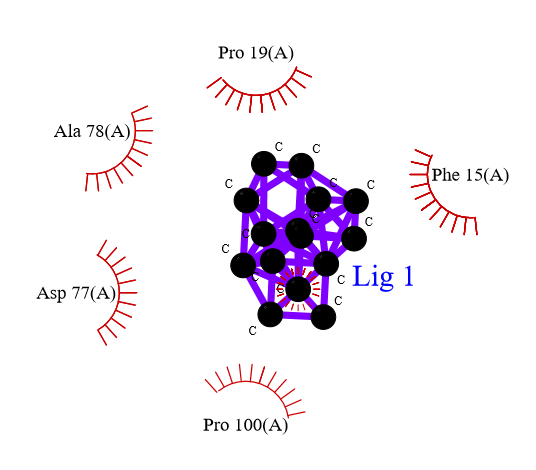

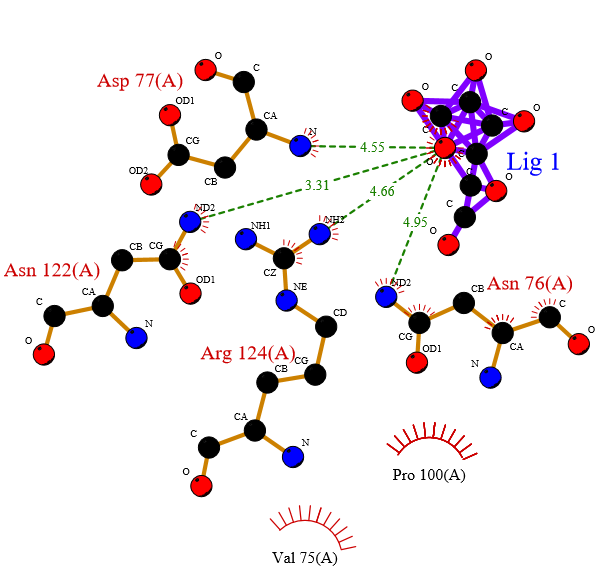

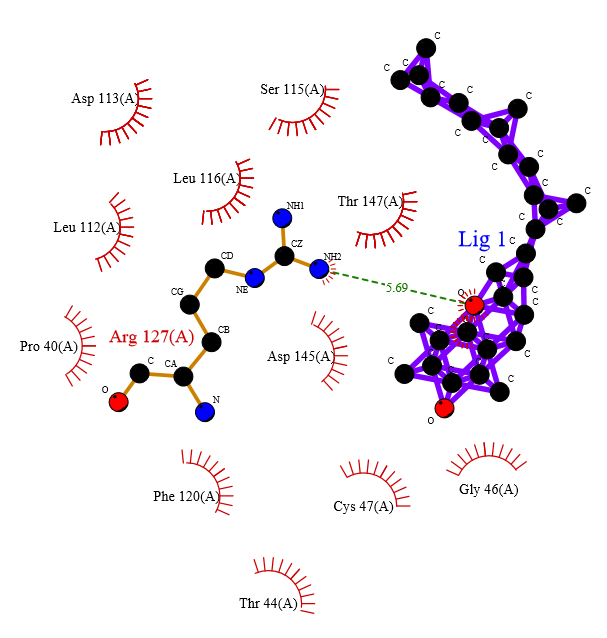

**Supplementary Fig. 2: LigPlot structure showing interactions between target protein (1HD2) with selected phytocompounds of CEO. 3,7-Nonadien-2-one, 8-methyl-, (E)- (A), α-pinene (B), Limonene (C), Citral (D), Epoxy- α -terpenyl acetate (E), Carane (F), 4,5-epoxy-, trans (G), 3-Cyclohexene-1-carboxaldehyde,1,3,4-trimethyl (H), Cyclohexane (I), Isopulegol (J), Camphene (J), Cis-verbenol (K), Caryophyllene oxide (L), Caryophyllene (M), Ascorbic acid (N), Tocopherol (O).**
